## Supplemental materials and methods for "*MYBPC3* D389V Variant Induces Hypercontractility in Cardiac Organoids"

### Expanded Materials and Methods

Running title: Cardiac Organoids for HCM

[€]Co-correspondence to Darshini Desai, MS, Division of Cardiovascular Health and Disease, University of Cincinnati, 231 Albert Sabin Way, Cincinnati, OH 45267, USA;


*Correspondence to Sakthivel Sadayappan, Ph.D., MBA, Division of Cardiovascular Health and Disease, University of Cincinnati, 231 Albert Sabin Way, Cincinnati, OH 45267, USA. Phone: +1 513 558 7498;

### iPSC Generation and Culture Conditions

Human subjects were enrolled under the Institutional Review Board (IRB) of University of Cincinnati guidelines (IRB# 2016-7580 and IRB# 2016-4948). All volunteers gave consent to the study. hiPSCs were generated at the Cincinnati Children's Pluripotent Stem Cell Facility from peripheral blood mononuclear cells (PBMCs) from Non-carrier and D389V carrier (age 52 and 54; 2 males). Thawed PBMCs were plated in erythroid expansion media (EEM; StemCell Technologies, Cat # 09860) and cultured for 8 days with regular addition of fresh EEM every 2 days. Following priming on day 0, cells were transduced with Sendai viral vectors (Cytotune 2.0, ThermoFisher Cat #A34546) via spinoculation and incubated overnight. On day 1, transduced cells were collected, resuspended, and further incubated for 48 hours. On day 3, transduced cells were co-cultured on gelatin-coated dishes with irradiated MEFs in complete StemPro-34 medium (Thermofisher, Cat #10639011). Medium changes were performed on days 4, 6, and 7, with a transition to hESC medium on day 7. Daily media changes with mTeSR™1 (Stem Cell Technologies, Cat # 85850) began on day 8. iPSC colonies were manually excised and replated in feeder-free culture conditions on matrigel-coated plates (Corning, Cat #CLS354277) in mTeSR1. Lines demonstrating robust proliferation and human pluripotent stem cells were expanded and cryopreserved at passage 8-12. hiPSCs were further cultured in mTeSR™1 plus supplement (Stem Cell Technologies, Cat. #85850) on a 6-well plate coated with hESC qualified Matrigel (Corning, Cat #CLS354277) in an incubator at 37 °C and 5% $CO_2$ until 80%-90% confluency was reached, at which point cells were split into 1:6 to 1:10 using Versene (Thermo Fisher Scientific, Cat #15040066) as a cell dissociation reagent.

### hiPSC Monolayer Cardiac Differentiation

Differentiation was performed using the small molecule Wnt pathway modulation approach adapted from the protocol of Lian et.al. [1] Briefly, hiPSCs at 80-90% confluency, are washed once with DPBS buffer (Thermo Fisher Scientific, Cat #14190144). Differentiation of hiPSCs to cardiomyocytes are generated in RPMI medium (Thermo Fisher Scientific Cat #11875093) with 2% B27 minus insulin supplement (Gibco Cat # A1895601) for 7 days of differentiation and then switched to RPMI with 2% B27 supplement (Gibco Cat #17504044) from days 7–30. On day 1, cells were treated with 5 µM CHIR99021 (STEM CELL TECHNOLOGIES, Cat #72052) for 24 h and with 2 µM IWP2 (STEM CELL TECHNOLOGIES, Cat #72122) for 48 h from day 3–5 of differentiation. After that, cells were incubated in RPMI/B27 minus insulin for two more days and then switched to RPMI/B27 complete medium until beating cardiomyocytes are observed. Beating CMs were maintained for up to 30 days in the RPMI/B27 medium which was changed every 48 hours.

### Generation of Human Cardiac Organoids using iPSC lines

After the dissociation using Versene, hiPSCs were centrifuged at 300 g for 5 min then resuspended in mTeSR™1 (Stem Cell Technologies, Cat # 85850) containing 1 µM ROCK inhibitor Y-27632 dihydrochloride (TOCRIS, Cat # 1254). Cells were then seeded at 250,000 cells in agarose molds on day −2 at a volume of 75 µl per mold (MicroTissues® 3D Petri Dish® micro-mold, Cat # Z764043). Each mold, containing 12X36 agarose wells, was placed in one well of a 12-well tissue culture plate. The plate was then centrifuged at 100 g for 3 min and placed in an incubator at 37 °C and 5% CO2. After 24 h (day 1), 2 ml of media was carefully removed from each well (containing mold), and 2ml of fresh mTeSR™1 medium was added. The plate was returned to the incubator for an additional 24 hours. On day 0, 1.5ml of medium was removed from each well and 1.5 ml of RPMI 1640/B-27, minus insulin (Gibco, Cat # A1895601) containing CHIR99021 (STEMCELL TECHNOLOGIES, Cat #72052) was added at a final concentration of 5µM/well along with BMP4(STEMCELL TECHNOLOGIES, Cat #78211) at 0.36 pM (1.25 ng/ml) and Activin A (STEMCELL TECHNOLOGIES, Cat #78001.1) at 0.08 pM (1 ng/ml) for 24 h. Next day, 1.5ml of media was removed and replaced with fresh RPMI1640/B-

27, minus insulin for 24h. On day 2, RPMI/B-27, minus insulin, containing IWP2 (STEM CELL TECHNOLOGIES, Cat #72122) was added to a final concentration of 5 μM and the samples were incubated for 48 h. The medium was changed again on day 4 with RPMI1640/B-27 minus insulin alone and cultured for another 48h. After Day 6, medium was switched to RPMI with B27 supplement (Gibco Cat #17504044) every 48hrs until organoids started beating and were ready for downstream analysis.

#### Confocal Microscopy and Image analysis

The cardiac organoids were characterized by staining for cardiac-specific markers. First, whole organoids were fixed in 4.0 % (v/v) paraformaldehyde/DPBS for 30 minutes. As organoids are like tissues, they were cleared using the CytoVista™ Tissue Clearing Kit (Invitrogen™, Cat #V11322). Organoids were then permeabilized through a series of incubations in increasing concentrations of methanol for 15 mins each (50%, 80%, 100%). Then the organoids were rehydrated through decreasing methanol concentrations (100%, 80%, 50%, PBS). Then the organoids were incubated in Antibody penetration buffer (provided in the kit). This was followed by incubation in blocking buffer for 1hr at 37 degrees Celsius. Organoids were then incubated overnight at 4 degrees Celsius with primary antibodies for cardiac and sarcomere markers as follows: anti-rabbit alpha-actinin-4 (Biorad, Cat #VPA00686, dilution 1:500), cMyBP-C; anti-mouse antibody (Santa Cruz Biotechnology, Cat # sc-137180, dilution 1:500), anti-mouse cardiac troponin I (Abcam, Cat # ab47003, dilution 1:500), Anti-rabbit Vimentin (Abcam, Catalog # ab16700, dilution 1:1000) in antibody dilution buffer provided with the kit. The next day, organoids were washed with PBS and wash buffer (provided with the kit )5 times (10mins each). After washing, the organoids were incubated with Goat Alexa Fluor 568 anti-rabbit and Goat Alexa Fluor 488 anti-mouse (Thermo Fisher Scientific Cat # A-11001 and A-11011, dilution 1:1000) secondary antibody for 1 hr at room temperature with gentle shaking. Afterwards, organoids were washed 10 times for 10 minutes with wash buffer. This was followed by dehydration with increasing methanol concentration (50%, 80% 100%). Organoids were then placed in the clearing reagent provided with the kit for 5 mins followed by mounting on a 30 cm dish with glass bottom (MatTek, Cat # P35G-1.5-20-C) using mounting medium containing DAPI to stain nuclei (SlowFade gold antifade Mountant, Thermo Fisher Scientific, Cat # S36936). A Leica dual confocal microscope was used to generate Z- stacks and 3D images at 10x, 20x and 63x magnification. Images were analyzed using Fiji (https://imagej.net/Fiji) Confocal microscopy support was provided by University of Cincinnati microscopy core.

#### Electron Microscopic Analyses

Organoids were fixed in 3% glutaraldehyde (Sigma Aldrich, Cat # G6257) in 0.15M sodium cacodylate buffer (Sigma Aldrich, Cat # 97068) postfixed in 1% osmium tetroxide (Sigma Aldrich, Cat # 20816-12-0) in 0.15M sodium cacodylate buffer, processed through a series of alcohols, infiltrated then embedded in LX-112 resin (LADD kit, Cat # 21210 - LX 112). After polymerization at 60 degrees for three days, ultrathin sections (120nm) were cut using a Leica EM UC7 ultramicrotome and counterstained in 2% aqueous uranyl acetate (VWR, Cat # 541-09-3) and Reynold's lead citrate (Sigma Aldrich, Cat # 15326). Images were taken with a transmission electron microscope (Hitachi H-7650) equipped with a digital camera (Biosprint 16). Electron microscopy support was provided by Integrated Pathology Research Facility at Cincinnati Children's Hospital.

#### Western Blot Analyses

To quantify expression levels of total and phosphorylated cMyBP-C, Western blot analysis was performed using organoid lysate. Thirty-forty organoids were pooled in 300 μl Protein Solubilization Buffer (PSB) (Biorad, Cat # 1632145) and 6-8 1.5 mm zirconia beads (Biospec products, Cat. # 11079101z) were added to each sample. Lysis was facilitated using a

BeadBlaster 24 (Benchmark Scientific) for 1 min at a speed of 7M/S followed by incubation for 10 min on ice. Cell debris was removed by centrifugation (12000 rpm, 10 mins, 4°C). Supernatants were transferred fresh tubes and protein concentrations were measured using the Bradford Plus protein assay reagent (Thermo Scientific, Pierce, Cat # 23238) according to the manufacturer´s instructions. For SDS-PAGE 30 µg protein, in 1X Laemmli Buffer plus 20mM DTT, was prepared for each sample and was then denatured for 5 min at 100°C.

Proteins were separated on a 4-20% Mini-Protean, TGX Stain-free gel (BioRad, Catalog # 4568093). Samples were run on three identical gels then transferred to nitrocellulose membranes (BioRad Cat. # 1620213). Transfer was performed on the benchtop packed in ice for 3 hrs at 250 mA in 1X Tris/Glycine Buffer plus 20% methanol. Membranes were cut horizontally between the 75 and 50kDa ladder bands then blocked with 5% dry milk (Blotting-grade blocker Non-Fat Dry Milk (BioRad, Cat. #1706404XTU) in TBS with 0.05% Tween20 (TBS-T) for 1 hour at RT. Following three washes with TBS-T, membranes were incubated overnight with primary antibodies diluted in TBS-T at 4°C. The top part of each membrane (≥75 kDa) was probed with one of three phospho-cMyBP-C antibodies (Rabbit anti-pSer275, pSer284 or pSer304 custom-made polyclonal antibody) at a dilution of 1:1000 [2]. The bottom part of each membrane (≤75 kDa) was probed with rabbit anti-GAPDH polyclonal antibody (Sigma-Aldrich, Catalog # G9545) at a dilution of 1:3000 [2]. This was followed by three washes with TBS-T. The membranes were incubated with Goat anti-rabbit IgG HRP-conjugated secondary antibody (Abcam, Catalog #. ab97080) prepared in 5% milk in TBS-T for 1 hr at RT. The protein of interest was then detected using Immobilon® Western Chemiluminescent HRP Substrate (Millipore, Cat. # WBKLS0500) and documented with a BioRad Chemidoc MP system. The phospho-cMyBP-C primary antibodies were stripped using the mild stripping protocol provided by AbCam. After ensuring that no residual signal was detected when re-probed with the secondary antibody, the membranes were probed with the cMyBP-C total protein antibody (Rabbit anti-cMyBP-C C2-14 custom-made polyclonal antibody) [3]. The same secondary antibody as before was used followed by chemiluminescent detection. Densitometric analysis was performed using ImageLab 6.0 (BioRad) software. The ratio of phosphorylated protein to total protein was calculated. To confirm equal loading of samples across the three blots cMyBP-C bands were normalized to the respective GAPDH values. Normalized expression values were plotted using GraphPad Prism 10 software.

### RNA Isolation, Sequencing and Transcriptomic analysis

2D cultured IPSC-CMs or pooled 3D cardiac organoids were washed with PBS and total RNA was isolated using the Rneasy Kit (Qiagen Cat # 74034) per the manufacturer's instructions. Lysis was performed in 350–500 µl Buffer RLT and RNA was eluted in 30-50 µl nuclease free water. RNA concentration was measured with a NanoDrop spectrophotometer. The RNA was subsequently stored at –80. Directional polyA RNA-seq was performed by the Genomics, Epigenomics and Sequencing Core at the University of Cincinnati as previously published [4, 5]. Quality control of the total RNA was performed using an Agilent 2100 Bioanalyzer (Agilent, Santa Clara, CA). PolyA RNA was isolated using the NEBNext Poly(A) mRNA Magnetic Isolation Module (New England BioLabs, Ipswich, MA) and 500 ng of total RNA input. The polyA RNA was then enriched using SMARTer Apollo automated NGS library prep system (Takara Bio USA, Mountain View, CA). The NEBNext Ultra II Directional RNA Library Prep kit (New England BioLabs, Cat. # E7760S) was used as a template for library preparation under PCR cycle number of 8. After which library quality control and quantitation was performed and using Qubit quantification (Thermo Fisher Scientific, Waltham, MA, Cat. # Q33221). Individually indexed libraries were evenly pooled and sequenced using a NextSeq 2000 Sequencer (Illumina, San Diego, CA) with the sequencing settings of PE 2x61 bp to produce approximately 60M reads.

After completion of the sequencing, fastq files for downstream data analysis were automatically generated via Illumina BaseSpace Sequence Hub. Paired-end reads were aligned against human hg19 genome using STAR (v2.6.1a, https://github.com/alexdobin/STAR). The raw gene counts were calculated using feature Counts (v1.5.2,http://subread.sourceforge.net/) and normalized using edgeR (v3.16.5, https://bioconductor.org/packages/release/bioc/html/edgeR.html).
Differentially expressed genes were predicted using limma/voom (v3.30.6, https://bioconductor.org/packages/release/bioc/html/limma.html). The default filtering cutoffs were set to (1) |fold| > 2x and (2) adjusted p-value < 0.05. Enrichment tests using up- and down-DEGs were performed against GOBP (biological process) and GOCC (cellular component) gene sets using EnrichR (https://maayanlab.cloud/Enrichr).

#### PCR primers and cDNA sequencing

2D cultured IPSC-CMs from NC and D389V cell lines were washed with PBS and total RNA isolation was performed using the Qiagen RNeasy kit according to the manufacturer´s instructions (Qiagen, Cat. # 74034). Cells were lysed in 350-500 µl of Buffer RLT and total RNA was eluted in 30 µl of nuclease free water. RNA was measured with a NanoDrop spectrophotometer (Thermo Scientific). 400 ng of total RNA was used to prepare cDNA using a Biorad iScript cDNA synthesis kit (BioRad, Cat # 1708890) according to the manufacturer's instructions and setup.

| **cDNA synthesis process** | **Time & temperature** |
|---|---|
| Priming | 5 min at 25°C |
| Reverse transcription | 20 min at 46°C |
| RT inactivation | 1 min at 95°C |
| Optional | Hold at 4°C |

The cDNA template was used to amplify the D389V mutation region (exon 12) of MYBPC3 using forward primer: 5'- GCA TGC TAA AGA GGC TCA AG - 3' and reverse primer: 5'- CTC CTC CGA TAC TTC ACA CTC A- 3' generating a 390bp amplicon.

| STEP | TEMP | TIME |
|---|---|---|
| Initial Denaturation | 95°C | 30 seconds |
| 30 Cycles | 95°C<br>45-47°C<br>68°C | 15-30 seconds<br>15-60 seconds<br>1 minute |
| Final Extension | 68°C | 5 minutes |
| Hold | 16°C | infinite |

The product was run on a 3% agarose gel for 1 hr at 120mV with a 100bp ladder (New England Biolabs, Cat # N3231L). The 390bp product was cut from the gel and extracted using the MinElute Gel Extraction Kit (Qiagen, Cat #28604) then sent for Sanger sequencing and analyzed using FinchTV.

### Spatial molecular imaging

Spatial molecular imaging was performed on organoid sections by NanoString Technologies Inc, Seattle, WA as described previously.6 Freshly isolated organoids were fixed in formalin and embedded in paraffin, sectioned at 4-6μm, and mounted on slides (VWR® Superfrost® Plus Micro Slide, Cat. # 48311-703). The slides were on a CosMx™ Spatial Molecular Imaging (SMI) platform (NanoString Technologies Inc, Seattle, WA), as described previously [6]. Briefly, slides were baked overnight at 60°C to ensure FFPE tissue adherence to the glass slides. Samples underwent deparaffinization, 3 μg/ml proteinase K digestion for 30 min at 40°C, and ER2 (Leica, AR9640) heat-induced epitope retrieval (HEIR) procedures to expose target RNAs and epitopes manually. The samples were fixed with 10% neutral buffered formalin (NBF) for 5 minutes at room temperature. Prepared samples were rinsed with 2X saline sodium citrate (SSC) for 5 min and then an Adhesive SecureSeal Hybridization Chamber (Grace Bio-Labs, Cat. # SKU: 621101) was placed to cover the samples.

CosMx Human RNA TAP Panel (1000-plex) plus a custom panel probe (*TNNI3, TNNT2, SERCA2, MYBPC3, MYH6, MYH7, NPPA*) mix were placed in the hybridization chamber overnight at 37°C. After hybridization, samples were washed twice with 50% formamide (VWR) in 2X SSC at 37°C for 25 min, rinsed twice with 2X SSC for 2 min at room temperature, then blocked with 100 mM NHS-acetate for 15 min. After blocking, the samples were washed twice using 2X SSC for 2 min at room temperature. Hybridization was performed at 37°C with this ISH probe mix followed by blocking with 100 mM NHS-acetate for 15 min. The flow cell was loaded onto the SMI instrument and scanned for 45 fields of view (FOVs) of one slide for RNA readout.

After RNA readout, organoid samples were incubated with a fluorophore-conjugated antibody cocktail against CD298 (Abcam, EP1845Y), B2M (Abcam, EP2978Y), PanCK (Novus, AE-1/AE-3), CD45 (Novus, 2B11 + PD7/26), and DAPI in the same instrument for 1 hr. Eight to nine Z-stack images for 5 channels (4 antibodies and DAPI) were captured.

The Z-stack images of nuclear (DAPI) and CD298/B2M (surface) staining were used to draw cell boundaries. A cell segmentation pipeline using a machine learning algorithm was used to accurately assign transcripts to cell locations and subcellular compartments. The transcript profile of individual cells was generated by combining target transcript location and cell segmentation boundaries. Cells with fewer than 20 total transcripts were omitted from downstream analysis. Expression profiles were normalized for each cell by dividing its raw count vector by total counts and multiplying by a scalar, the average total counts per cell. Cells were clustered using unsupervised clustering as implemented in the InSituType package. Clustered cells were displayed using the runUMAP function in Giotto. Differential expression tests modeled count data with a negative binomial distribution and included the expression of genes in neighboring cells as a fixed effect to reduce the impacts of contamination due to minor mis-segmentation.

### hCOs Contractility and Calcium Handling

Cardiac contractility was assessed on spontaneously beating cardiac organoids in a 30mm culture dish using the CytoCypher MultiCell High Throughput System (CytoCypher BV, IonOptix) [7]. Cardiac organoid contractile kinetics were measured at 37°C and based on brightfield CytoMotion pixel intensity changes relative to a diastolic reference frame acquired at 250Hz sampling frequency (IonOptix LLC). Simultaneously, calcium transients were measured by loading organoids with 0.5μM Fura-2/AM (Thermo Fisher Scientific, Cat # F1221) for 25 min in RPMI/B27 complete medium, followed by medium wash. We measured 25 to 60 organoids for 10s, in which 4-10 contraction traces were recorded with or without Mavacamten (MYK-461,

MCE, Cat #HY-109037) in the medium. The CytoSolver Transient Analysis Tool package from CytoCypher BV/IonOptix was used to yield averaged contractile and calcium kinetic parameters from each area. For the purposes of this study, four key parameters of contractility were analyzed: contraction time or time to peak 50% and 90% (s) and relaxation time or time to baseline 50% and 90% (s). For calcium, we analyzed six key parameters, time to peak 50% and 90%, time to decay 50% and 90% and calcium amplitude and baseline calcium levels (diastolic calcium).

#### Quantification of ROS generation and Membrane potential

ROS generation was measured in live hCOs using the Reactive Oxygen Species (ROS) Detection Assay Kit (Abcam, Cat # ab287839) according to the manufacturer's instructions. The hCOs from both the genotypes were incubated with H2DCFDA dye for 30mins with and without N-acetyl cysteine (NAC, antioxidant as a negative control). Oxidation of H2DCF by intracellular ROS highly produces a fluorescent product that is detected on a fluorescence microscope (Ex/Em 495/529 nm) [8]. Image J to calculate the intensity in each organoids using the integrated intensity tool. Background fluorescent intensities were subtracted, and resultant values were plotted for to compare D389V and NC hCOs. The following formula was used to measure the intensity: Corrected total cell fluorescence (CTCF). = Integrated Density – (Area of selected cell X Mean fluorescence of background readings).

The mitochondrial membrane potential was measured by fluorescence detection using the JC-10 Assay Kit (Abcam, Cat # ab112134) according to the manufacturer's instructions. The cells were isolated from hCOs by incubating them in warm TrypLE for 10mins and resuspended single cells into warm RPMI/B27 complete medium. The single hiPSC-CM were then plated on Matrigel coated 96-well, black-walled, clear-bottom plates and cultured for 4-5 days, until they start beating. Cells were then stained with 50 mL of JC-10 solution. The fluorescence intensity (excitation/emission [Ex/Em] = 485/525 nm and Ex/Em = 540/590 nm) was measured on a Synergy H1 Hybrid microplate reader (BioTek).

#### Co-sedimentation Assay

To determine whether D389V mutation in the C2 domain affects the cMyBP-C- Myosin S2 affinity, a co-sedimentation assay was performed as described previously [9]. For this assay, we generated recombinant human wild-type (WT) and D389V C0-C2 proteins as described previously [9]. These proteins are approximately 65 kilodaltons in size. Recombinant C0-C2 proteins were then dialyzed against co-sedimentation buffer (Identical to high-salt myosin storage buffer, but without KCl) overnight in the cold room and concentrations were determined by Bradford assay. Varying concentrations of C0-C2 proteins (0.1 – 4μM) were mixed with a fixed amount of myosin and brought to a final volume of 50μl with a final KCl concentration of 100mM and final myosin concentration of 1μM. Samples were incubated for 30 minutes at room temperature with agitation and then centrifuged for 1 hour at 178,000 x g. Pellets were resuspended in sample buffer and loaded onto 10% SDS-PAGE gels. Standards were prepared similarly but without centrifugation, supplementing co-sedimentation buffer with sample buffer. Gels were stained with Coomassie blue, and the intensity ratio of C0-C2 to myosin was converted to a mole/mole ratio as previously described [10]. Curves were fit to a one-site binding (hyperbola) to report $B_{max}$ and $K_d$.

#### Solid-phase Binding Assays (SPBA)

SPBA was measured using the previously described protocol with minimal modifications [10]. Briefly, 20μM of C0C2 peptides were adsorbed onto the 96-well plates then incubated with 3% skim milk to block nonspecific binding. Proximal myosin S2 peptides were added in increasing concentrations (0-20 μM) and incubated overnight at 4°C. Bound myosin S2 was detected using a rabbit polyclonal primary antibody and Alexa Fluor-568 goat anti-rabbit as the secondary antibody. The plates were read using the Cytation™ 5 Cell Imaging Multi-Mode Reader (BioTek) with excitation at 568 nm and emission at 619 nm. Binding curves were plotted in GraphPad Prism 9.0 and fit using Michaelis-Mentin nonlinear regression to obtain relative Vmax and km values.

### Isothermal calorimetry assay (ITC)

To validate the SPBA experiments, ITC experiments were conducted to determine the interactions between hS2 and hC0-C2$^{WT}$, and hS2 and hC0-C2$^{D389V}$ as described previously using the MicroCal VP-ITC 2000 calorimeter [10]. Briefly, 20 µM of hC0-C2$^{WT}$ and hC0-C2$^{D389V}$ protein was titrated against 350 µM of S2 protein in PBS at 20 °C. Before titration, the protein solutions were degassed. To elucidate the true thermal interaction between recombinant hS2 and hC0-C2 proteins, diluent and titrant controls were used. In each experiment, a total of 25 injections of titrant, each consisting of ten microliters, were administered at intervals of 180 seconds. Next, a titration speed of 351 rpm was implemented to ensure binding. ΔH, Kd, and η were calculated by fitting the amount of heat produced at each titration and the η of hS2 and hC0-C2 to a four-parameter logistic curve (4PL) Y = y0 + (xˆHillslope)∗(Top-Bottom)/(xˆHillSlope+EC50ˆHillSlope) model in GraphPad Prism 9.

### *In vitro* Motility Assay

To measure the function of actin and myosin filaments in the presence of D389V mutation, we performed *in vitro* motility assay using actin filaments sliding over myosin filaments on a 2% dimethyldichlorosilane-coated coverslip as described previously [11]. The suspension contained 1mM ATP diluted in 4 mM magnesium chloride, 0.5 mg/ml bovine serum albumin, 10 mM imidazole, 10 mM dithioerythritol, 0.5% methylcellulose, 1% glucose, 45 µg/ml catalase 25 mM, potassium chloride, and 25 µg/ml glucose oxidase at 30 °C. Bovine Myosin HMM[12] and F-actin (Thermo Fisher Scientific, Cat # R415) filaments, were labeled with rhodamine-phalloidin and excited with a green LED light source on an epifluorescent microscope and imaged with a TIRF microscope (Nikon A1R LUN-V inverted confocal fluorescent microscope). The microscope stage heater, maintained the temperature at 30 °C. The motility of actin was measured before and after adding an equimolar concentration of C0-C2 recombinant proteins to the myosin HMM. Images were then analyzed by Image J (NIH) software. The manual tracking Image J plugin was used to record the movements of actin filaments.

### Statistics and Reproducibility

All analyses were performed using GraphPad software and all raw data was collected in Microsoft Excel and were normally distributed. Statistical significance was evaluated with a standard unpaired Student t-test (2-tailed; $P < 0.05$) when appropriate (Western blot, ROS generation, Membrane potential experiments). For binding and in-vitro motility experiments one-way ANOVA with Tukey's multiple comparison test with single pooled variance ($n = 3$, $P < 0.05$) was used. For multiple-comparison analysis, 2-way ANOVA with post-test correction applied ($P < 0.05$). All data are presented as mean ± SEM.
